# Genetic determinants of lipids and cardiovascular disease outcomes: a wide-angled Mendelian randomization investigation

**DOI:** 10.1101/668970

**Authors:** Elias Allara, Gabriele Morani, Paul Carter, Apostolos Gkatzionis, Verena Zuber, Christopher N Foley, Jessica MB Rees, Amy M Mason, Steven Bell, Dipender Gill, Adam S Butterworth, Emanuele Di Angelantonio, James Peters, Stephen Burgess

## Abstract

**Aims:** To systematically investigate causal relationships between circulating lipids and cardiovascular outcomes, using a Mendelian randomization approach.

**Methods and Results:** In the primary analysis, we performed two-sample multivariable Mendelian randomization using data from participants of European ancestry. We also conducted univariable analyses using inverse-variance weighted and robust methods, and gene-specific analyses using variants that can be considered as proxies for specific lipid-lowering medications. We obtained associations with lipid fractions from the Global Lipids Genetics Consortium, a meta-analysis of 188,577 participants, and genetic associations with cardiovascular outcomes from 367,703 participants in UK Biobank.

For LDL-cholesterol, in addition to the expected positive associations with coronary artery disease (CAD) risk (odds ratio per 1 standard deviation increase [OR], 1.45; 95% confidence interval [95%CI] 1.35-1.57) and other atheromatous outcomes (ischemic cerebrovascular disease and peripheral vascular disease), we found independent associations of genetically-predicted LDL-cholesterol with abdominal aortic aneurysm (OR 1.75; 95%CI 1.40-2.17), and aortic valve stenosis (OR 1.46; 95%CI 1.25-1.70). Genetically-predicted triglyceride levels were positively associated with CAD (OR 1.25; 95%CI 1.12-1.40), aortic valve stenosis (OR 1.29; 95%CI 1.04-1.61), and hypertension (OR 1.17; 95%CI 1.07-1.27), but inversely associated with venous thromboembolism (OR 0.79; 95%CI 0.67-0.93). The positive associations of genetically-predicted LDL-cholesterol and triglycerides with heart failure and aortic stenosis appeared to be mediated by CAD.

**Conclusion:** Lowering LDL-cholesterol is likely to prevent abdominal aortic aneurysm and aortic stenosis, in addition to CAD and other atheromatous cardiovascular outcomes. Lowering triglycerides is likely to prevent CAD and aortic valve stenosis, but may increase risk of thromboembolism.

## INTRODUCTION

Evidence from randomized trials has shown that therapies that lower low-density lipoprotein (LDL)-cholesterol, such as statins, are beneficial for preventing or treating several atheromatous diseases such as coronary artery disease (CAD),^1^ ischaemic stroke,^2^ and peripheral vascular disease,^3,4^ as well as both postoperative atrial fibrillation^5^ and heart failure in the presence of underlying CAD.^6,7^ However, for other cardiovascular outcomes, such as thromboembolic disease,^8^ haemorrhagic stroke^9,10^ and aortic aneurysms,^11,12^ the effects of LDL-cholesterol lowering are less clear. Recently, the REDUCE-IT trial has shown that therapies that predominantly lower triglycerides can reduce major cardiovascular events.^13^ However, the effects of triglyceride lowering on non-atheromatous cardiovascular outcomes are largely unknown.

Although a randomized trial is the gold standard of evidence and is required to conclusively establish the effectiveness of a treatment, naturally-occurring genetic variants can be used to help predict the outcome of a randomized trial in an approach known as Mendelian randomization.^14^ For example, this approach has been used successfully to validate the effects of statins and predict the effects of PCSK9 inhibitors on CAD risk seen in randomized trials.^15,16^ Mendelian randomization investigations have also shown positive associations of genetically-predicted LDL-cholesterol with abdominal aortic aneurysm^17^, ischaemic stroke,^18^ and aortic stenosis,^19^ and positive associations of triglycerides with CAD.^20^

However, Mendelian randomization approaches have not yet been used to investigate the relationship between circulating lipids and some major cardiovascular conditions such as venous thromboembolism and heart failure. Additionally, approaches to estimate the independent effects of LDL-cholesterol, triglycerides, and high-density lipoprotein (HDL)-cholesterol on cardiovascular disease while accounting for genetic pleiotropy have only been performed for a limited set of outcomes.^17,19,20^

Our objective in this paper is to assess which cardiovascular outcomes could be treated by lipid-lowering therapies. We performed a systematic Mendelian randomization analysis of atheromatous and non-atheromatous cardiovascular disease outcomes. We considered disease outcomes in a single dataset (UK Biobank) to ensure a consistent approach to the analysis across different outcomes. We carried out polygenic analyses for each of LDL-cholesterol, HDL-cholesterol, and triglycerides, based on all common genetic variants associated with at least one of these risk factors, as well as gene-specific analyses based on variants in or near gene regions that mimic specific pharmaceutical interventions.

## METHODS

### Study design and data sources

We performed two-sample Mendelian randomization analyses, taking genetic associations with lipid concentrations from one dataset, and genetic associations with cardiovascular disease outcomes from an independent dataset.

We obtained genetic associations with lipid concentrations (LDL-cholesterol, HDL-cholesterol, and triglycerides) from the Global Lipids Genetics Consortium (GLGC) on up to 188,577 participants of European ancestry.^21^ We estimated genetic associations with disease outcomes on 367,703 unrelated participants of European ancestry from UK Biobank, a population-based cohort recruited between 2006-2010 at 22 assessment centres throughout the UK and followed-up until 31^st^ March 2017 or their date of death (recorded until 14^th^ February 2018).^22^ We defined 19 outcomes based on electronic health records (ICD-9 or ICD-10 diagnosis), hospital procedure codes, and self-reported information validated by interview with a trained nurse (Supplementary Table S1). We included CAD as a positive control outcome, and chronic kidney disease as a negative control outcome as it is thought to be influenced by some cardiovascular risk factors (such as hypertension and diabetes) but not dyslipidaemia.^23^

### Statistical analysis

We carried out polygenic analyses based on 184 genetic variants previously demonstrated to be associated with at least one of LDL-cholesterol, HDL-cholesterol or triglycerides at a genome-wide level of significance (*p* < 5 × 10^−8^) in the GLGC. These variants explained 13.7% of the variance in HDL-cholesterol, 14.6% in LDL-cholesterol, and 11.7% in triglycerides in GLGC.

To obtain the associations of genetically-predicted values of these lipids with each cardiovascular outcome while accounting for measured genetic pleiotropy via other major lipids, we performed multivariable Mendelian randomization analyses.^24^ As sensitivity analyses, we also performed (i) univariable Mendelian randomization using the inverse-variance weighted method for each lipid fraction based on all variants associated with that lipid fraction at a genome-wide level of significance,^25^ (ii) MR-Egger regression to account for unmeasured pleiotropy,^26^ and (iii) weighted median regression to assess robustness to invalid genetic instruments.^27^ To account for between-variant heterogeneity, we used random-effects models in all analyses. All Mendelian randomization estimates are expressed per 1 standard deviation increase in the corresponding lipid fraction in GLGC (1 standard deviation was 39.0 mg/dL for LDL-cholesterol, 15.8 mg/dL for HDL-cholesterol, and 90.5 mg/dL for triglycerides).

For cardiovascular diseases for which it is possible that effects of lipid fractions on that outcome may act via other conditions (e.g. via CAD or type 2 diabetes), we also performed multivariable Mendelian randomization analyses adjusted for the possible mediating condition(s). To validate results for thromboembolism outcomes, we performed additional analyses for traits related to thrombosis and coagulation that have publicly-available genetic association data.^28^

We also performed gene-specific analyses for variants in gene regions that can be considered as proxies for existing or proposed lipid-lowering therapies. We conducted separate analyses for the *HMGCR, PCSK9, LDLR, APOC3*, and *LPL* gene regions (Supplementary Table S2). These regions were chosen as they contain variants that explain enough variance in lipids to perform adequately powered analyses.

Further information on the study design and statistical analysis is available in the Supplementary Methods.

## RESULTS

### Participant characteristics

Baseline characteristics of the participants in the UK Biobank are provided in Table 1. Around 46% of participants were men, and the mean age was 57 years. Around 10% were smokers, 93% were alcohol drinkers, and 4% had a history of diabetes at baseline.

**Table 1:** Baseline characteristics of UK Biobank participants included in this study and numbers of outcome events

| Characteristic | Mean (SD) or N (%) |
| --- | --- |
| Sample size | 367,703 (100) |
| Male | 168,799 (45.9) |
| Age at baseline | 57.2 (8.1) |
| Body mass index | 27.3 (4.8) |
| Systolic blood pressure | 137.6 (18.6) |
| Diastolic blood pressure | 82.0 (10.1) |
| Smoking status (current / ex / never)* | 37,866 (10.3) / 185,704 (50.5) / 143,777 (39.1) |
| Alcohol status (current / ex / never)* | 342,797 (93.2) / 12,732 (3.5) / 11,646 (3.2) |
| History of type 2 diabetes | 15,834 (4.3) |
| Coronary Artery Disease | 29,278 (8.0) |
| Ischaemic Cerebrovascular Event (all) | 8084 (2.2) |
| - Ischaemic Stroke | 4602 (1.3) |
| - Transient Ischaemic Attack | 3962 (1.1) |
| Haemorrhagic Stroke (all) | 1981 (0.5) |
| - Intracerebral Haemorrhage | 1064 (0.3) |
| - Subarachnoid Haemorrhage | 1084 (0.3) |
| Aortic Aneurysm (all) | 1849 (0.5) |
| - Abdominal Aortic Aneurysm | 1094 (0.3) |
| - Thoracic Aortic Aneurysm | 347 (0.1) |
| Venous Thromboembolism (all) | 14,097 (3.8) |
| - Deep Vein Thrombosis | 9454 (2.6) |
| - Pulmonary Embolism | 6148 (1.7) |
| Hypertension | 125,846 (34.2) |
| Peripheral Vascular Disease | 3415 (0.9) |
| Aortic Valve Stenosis | 2244 (0.6) |
| Atrial Fibrillation | 16,945 (4.6) |
| Heart Failure | 6712 (1.8) |
| Chronic Kidney Disease | 6321 (1.7) |

\* Excluding 356 participants with smoking status absent and 528 participants with alcohol consumption status absent

### Polygenic analyses for all lipid-related variants

Multivariable Mendelian randomization estimates are displayed graphically in Figure 1 and summarized in Supplementary Table S3. Associations with the positive control outcome (CAD) were as expected for LDL-cholesterol and triglycerides. There were no associations with the negative control outcome (chronic kidney disease).

**Figure 1:**
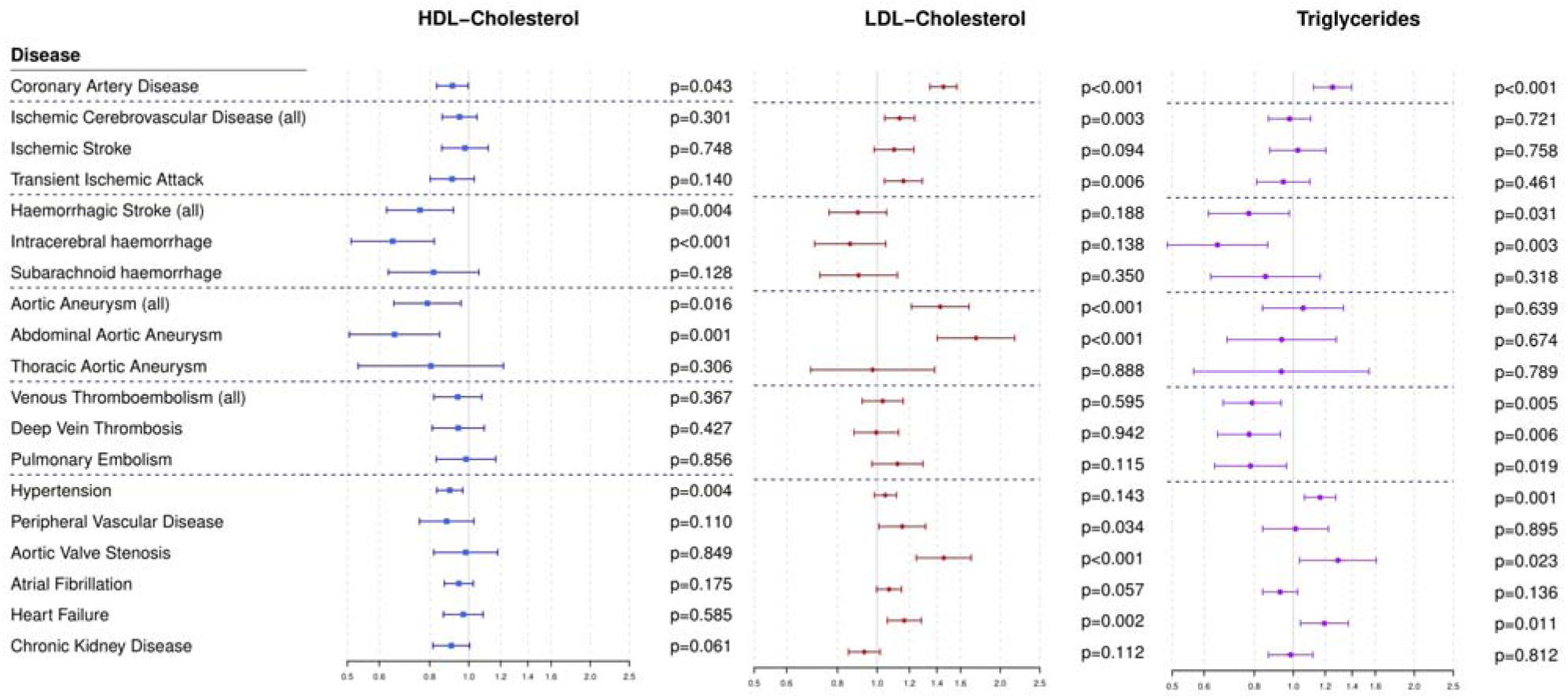
Multivariable Mendelian randomization estimates (odds ratio with 95% confidence interval per 1 standard deviation increase in lipid fraction) from polygenic analyses including all lipid-associated variants

We found a strong association of genetically-predicted LDL-cholesterol with CAD risk (odds ratio per 1 standard deviation increase [OR], 1.45; 95% confidence interval [CI] 1.35-1.57), which was in the reference range of a previous study.^20^ We also saw positive associations for LDL-cholesterol with risk of transient ischaemic attack, peripheral vascular disease, aortic aneurysm, abdominal aortic aneurysm and aortic valve stenosis. Of note, we found a novel positive association with heart failure (OR 1.17; 95%CI 1.06-1.28). The strongest associations by magnitude were for abdominal aortic aneurysm (OR 1.75; 95%CI 1.40-2.17) and aortic valve stenosis (OR 1.46; 95% CI 1.25-1.70). We observed a positive association for the combined outcome of ischaemic stroke and transient ischaemic attack (OR 1.14; 95% CI 1.04-1.24). The association with ischaemic stroke was in the positive direction but non-significant (OR 1.10; 95% CI 0.98-1.23).

Genetically-predicted triglyceride levels were associated with increased risk of CAD, consistent with previous results,^20^ as well as with increased risk of hypertension, aortic valve stenosis, and heart failure. Inverse associations were observed for triglycerides with all of the thromboembolic diseases we analysed: deep vein thrombosis (OR 0.78; 95%CI 0.65-0.93), pulmonary embolism (OR 0.78; 95%CI 0.64-0.96), and any venous thromboembolism (OR 0.79; 95%CI 0.67-0.93). An inverse association was observed with haemorrhagic stroke, in particular with intracerebral haemorrhage (OR 0.65; 95%CI 0.49-0.86). Additionally, genetically-predicted triglyceride levels were associated with several traits related to thrombosis and coagulation, including red cell distribution width and tissue-type plasminogen activator, with the general direction of associations corresponding to an anti-thrombotic effect of increased triglycerides (Supplementary Table S4). Genetically-predicted HDL-cholesterol was weakly associated with coronary heart disease (OR 0.91; 95%CI 0.83-1.00), and more strongly inversely associated with abdominal aortic aneurysm, hypertension and haemorrhagic stroke, in particular intracerebral haemorrhage (OR 0.65; 95%CI 0.51-0.82).

Univariable Mendelian randomization estimates for each lipid risk factor in turn are displayed in Supplementary Figures S1-S3 and summarized in Supplementary Tables S5-S7. Similar results were generally obtained from each of the univariable methods as for the multivariable methods, although confidence intervals were slightly wider in some cases, especially for the MR-Egger method. The association between HDL-cholesterol and CAD became null in MR-Egger regression (OR 0.99; 95%CI 0.83-1.19). Similarly, the associations of triglycerides with haemorrhagic stroke and its subtypes attenuated to the null in almost all univariable analyses (Supplementary Table S7). Another notable difference was a positive association of increased genetically-predicted triglycerides with aortic aneurysm risk, particularly for abdominal aortic aneurysm. While the association with aortic aneurysm was not strongly apparent in the multivariable analysis, it was evident for each of the univariable analysis methods.

We also assessed whether the associations of genetically-predicted lipids with outcomes that are comorbid with CAD (e.g. heart failure) may be mediated via CAD, and similarly for outcomes that are comorbid with type 2 diabetes (e.g. hypertension). On adjustment for CAD risk (Table 2), the associations of genetically-predicted LDL-cholesterol and triglycerides with heart failure attenuated sharply, suggesting that their effects on heart failure might be mediated via CAD. As a negative control, we also performed the same analysis for abdominal aortic aneurysm. The association with genetically-predicted LDL-cholesterol attenuated slightly, but was still clearly positive. We also performed multivariable Mendelian randomization analyses excluding participants with CAD for heart failure (2383 remaining cases). This analysis gave null estimates for all lipid fractions (Supplementary Table S8). This suggests that LDL-cholesterol is unlikely to be a causal risk factor for heart failure where there is no comorbidity with CAD. On adjustment for body mass index (BMI) and type 2 diabetes (Supplementary Table S9), associations between lipids and hypertension, abdominal aortic aneurysm, and haemorrhagic stroke did not change substantially compared to unadjusted analyses, suggesting that neither BMI nor type 2 diabetes is likely to mediate the effects of lipids on these outcomes.

**Table 2:** Multivariable Mendelian randomization estimates (odds ratio with 95% confidence interval) for heart failure without and with adjustment for coronary artery disease (CAD). Analyses were also performed for abdominal aortic aneurysm as a negative control.

| ***Heart failure*** | **HDL-cholesterol** | **LDL-cholesterol** | **Triglycerides** | **CAD** |
| --- | --- | --- | --- | --- |
| *Without adjustment* | 0.97<br>(0.87-1.08) | 1.17<br>(1.06-1.28)** | 1.19<br>(1.04-1.37)* | - |
| *With adjustment* | 1.02<br>(0.92-1.13) | 0.95<br>(0.85-1.06) | 1.06<br>(0.93-1.20) | 1.73<br>(1.47-2.04)*** |
| ***Abdominal aortic aneurysm*** | **HDL-cholesterol** | **LDL-cholesterol** | **Triglycerides** | **CAD** |
| *Without adjustment* | 0.65<br>(0.51-0.85)** | 1.75<br>(1.40-2.17)** | 0.94<br>(0.68-1.28) | - |
| *With adjustment* | 0.69<br>(0.53-0.88)** | 1.46<br>(1.12-1.91)** | 0.84<br>(0.61-1.16) | 1.60<br>(1.06-2.42)* |

\*: $p < 0.05$ \*\*: $p < 0.005$ \*\*\*: $p < 10^{-8}$
The CAD column indicates the association between genetically-predicted coronary artery disease and the outcome of interest (i.e. heart failure or abdominal aortic aneurysm) while accounting for the genetic associations of the lipid fractions in a multivariable analysis. If the regression coefficients for the lipid measurements remain unchanged on adjustment for CAD, then the effects of lipids on the outcome do not operate via CAD. If the regression coefficients attenuate to the null, then the effects of lipids on the outcome are entirely mediated via CAD.

### Gene-specific analyses for drug proxy variants

Mendelian randomization estimates for specific gene regions are displayed graphically in Figure 2 and summarized in Supplementary Tables S10-S11. All the gene regions analysed showed clear associations with CAD risk, confirming their involvement in cardiovascular disease aetiology and the relevance of existing and proposed lipid-lowering therapies for CAD risk reduction.

**Figure 2:**
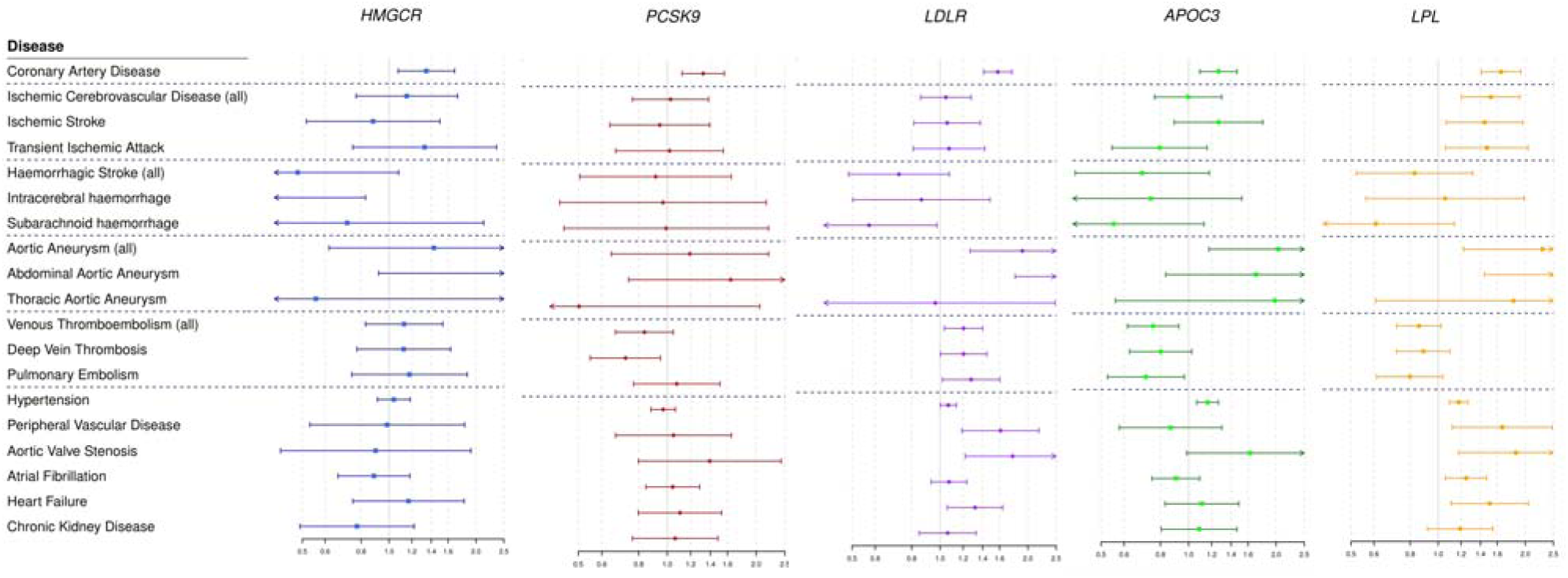
Univariable Mendelian randomization estimates (odds ratio with 95% confidence interval per 1 standard deviation increase in lipid fraction) for variants in specific gene regions. Estimates are scaled to a unit standard deviation increase in LDL-cholesterol for the *HMGCR, PCSK9*, and *LDLR* regions, and to a standard deviation increase in triglycerides for the *APOC3* and *LPL* regions.

Results for the *LDLR* gene regions, and to a lesser extent for *HMGCR* and *PCSK9*, were similar to those for LDL-cholesterol in the polygenic analyses, although with wider confidence intervals. Significant positive associations were obtained for *LDLR* with aortic and abdominal aortic aneurysm, venous thromboembolism, aortic valve stenosis, and heart failure, and an inverse association was observed for subarachnoid haemorrhage. Variants in the *HMGCR* gene region were inversely associated with intracerebral haemorrhage (*p*=0.022). Results for *APOC3* and *LPL* followed a pattern more similar to the analysis for triglycerides, showing inverse associations with thromboembolic diseases (*p*=0.007 for *APOC3, p*=0.089 for *LPL* for any venous thromboembolism). In both regions, associations with aortic aneurysm, aortic valve stenosis and hypertension were all positive. Additionally, variants in the *LPL* region were also positively associated with all ischaemic cerebrovascular diseases.

## DISCUSSION

In this study, we assessed the causal role of three major lipid fractions for a range of cardiovascular diseases in a large population-based cohort using the principle of Mendelian randomization while accounting for genetic pleiotropy between the lipid measures. Our most notable findings were the associations between genetically-predicted triglycerides and decreased risk of thromboembolic diseases both in polygenic analyses and in gene-specific analyses for the *APOC3* gene region, suggesting that reducing triglycerides may increase risk for venous thromboembolism. Additionally, we found: 1) evidence supporting the current understanding of the aetiology of CAD, suggesting independent causal roles for LDL-cholesterol and triglycerides, both in the polygenic analyses and for all the drug-related gene-specific analyses^20,29^ 2) positive associations of genetically-predicted LDL-cholesterol with abdominal aortic aneurysm and aortic valve stenosis, as well as atheromatous cardiovascular outcomes that are already addressed in clinical guidelines (e.g. peripheral vascular disease and the combined outcome of ischaemic stroke and transient ischaemic attack); 3) positive associations of genetically-predicted LDL-cholesterol and triglycerides with heart failure that appear to be mediated by CAD; 4) associations between genetically-predicted triglycerides and increased risk of aortic stenosis and hypertension; and 5) inverse associations between genetically-predicted HDL-cholesterol and abdominal aortic aneurysm, hypertension, and haemorrhagic stroke (in particular, intracerebral haemorrhage) that appear to be independent of BMI/T2D.

The importance of these findings is threefold. Firstly, they help us to better understand the aetiology and pathophysiology of common cardiovascular outcomes, and so to identify additional potential indications for lipid-lowering therapies. Secondly, association estimates provided in this paper can inform calculations on the risk-benefit and cost-benefit of these therapies. Thirdly, these findings can aid identification of which patients would most benefit from (or should avoid) lipid-lowering therapies.

Genetic predisposition to lower triglycerides was associated with an increased risk of thromboembolic diseases. In contrast, previous observational evidence did not suggest an association between triglycerides and venous thrombosis,^30,31^ although other lipid measures (in particular apolipoprotein B and lipoprotein(a)) were inversely associated with venous thrombosis mortality in a meta-analysis of over 700,000 participants from the Emerging Risk Factors Collaboration.^31^ Our result was consistent across 3 of the 4 Mendelian randomization methods. However, associations with thromboembolic outcomes using the MR-Egger method were slightly attenuated and compatible with the null, raising the possibility of unmeasured genetic pleiotropy. Variants in the *APOC3* gene region were associated with thromboembolic events, suggesting that lowering triglyceride levels via intervening on this pathway may increase thromboembolism risk. Additionally, associations between genetically-predicted triglycerides and molecular traits relating to thrombosis, coagulation and fibrinolysis were generally concordant with this finding, providing some validation of this claim. Further investigation is needed to confirm this unexpected finding.

Both the positive association between genetic predictors of LDL-cholesterol and abdominal aortic aneurysm, and the inverse association for HDL-cholesterol, replicate a previous Mendelian randomization study which did not include UK Biobank participants.^17^ This evidence is complemented by a recent randomized trial that has shown benefit from screening and treating abdominal aortic aneurysm using various interventions including statins,^11^ by observational studies that have shown positive associations between LDL-cholesterol and abdominal aneurysm,^32,33^ as well as an animal study demonstrating that elevated LDL-cholesterol levels led to atherogenesis, abdominal aneurysm formation, and increased mortality in a murine model.^34^ However, no significant association was seen with thoracic aortic aneurysm. Discrepancies between the LDL-cholesterol estimates for abdominal and thoracic aortic aneurysm (*p*=0.006 for difference) are in line with the different pathophysiology of the two disease subtypes.^35^ Overall, these findings provide further support to the hypothesis that increased LDL-cholesterol has deleterious effects on abdominal aortic aneurysm, and suggest that LDL-cholesterol lowering therapies may be beneficial in preventing abdominal aortic aneurysm.

Similarly, the positive association between genetically-predicted LDL-cholesterol and aortic valve stenosis replicates a previous Mendelian randomization study.^19^ We also demonstrated a positive association for triglycerides, in line with the previous study (although their result did not achieve conventional levels of statistical significance). Our association was robust to all sensitivity analyses and evidenced in gene-specific analyses for both the *APOC3* and *LPL* loci. These findings are consistent with previous observational and pathological studies suggesting a role of atherosclerotic processes in early valve lesions^36,37^ whereas clinical trials showed no benefit from LDL-cholesterol lowering via statins on aortic stenosis progression.^38^ Taken together, this suggests that increased LDL-cholesterol and triglycerides may facilitate initiation of early lesions, but that lipid-lowering therapies may be ineffective in preventing the progression of aortic stenosis.

CAD is a well-known risk factor for heart failure,^39^ as myocardial ischaemic damage reduces myocardial contractility and ventricular function. Associations of heart failure with genetically-predicted LDL-cholesterol and triglycerides disappeared after adjusting for CAD. The association was also absent when omitting participants with a CAD diagnosis from analyses. In contrast, associations with abdominal aortic aneurysm attenuated only slightly after adjustment for CAD. Overall, these results suggest that lipid-lowering therapies are only likely to influence risk of heart failure via their effects on CAD.

Our analyses also identified inverse associations of genetically-predicted HDL-C and triglyceride levels with risk of intracerebral haemorrhage, consistent with a recent observational analysis for triglycerides.^40^ Furthermore, we identified a detrimental effect of HMGCR inhibition on intracerebral haemorrhage risk, in keeping with a large randomized trial of atorvastatin in patients with recent transient ischaemic attack or stroke, which found statin therapy to lower risk of major cardiovascular events but increase the risk of haemorrhagic stroke.^41^ However, this finding has not been replicated in large meta-analyses.^42^ The mechanism relating lipid traits with risk of intracerebral haemorrhage requires further exploration.

Our investigation has both strengths and limitations. The large sample size of over 360,000 participants and the broad set of outcomes analysed render this one of the largest and most comprehensive Mendelian randomization analyses on cardiovascular disease conducted to date. Availability of multiple cardiovascular conditions within the same study enabled cross-comparisons between diseases, making this the first Mendelian randomization investigation to systematically assess the independent effects of circulating lipids on such a broad set of cardiovascular outcomes. This also enabled us to perform adjusted analyses in the same participants, to assess mediation of causal effects by CAD and other potential mediators. This investigation has a number of limitations. Lack of publicly-available summary data from external datasets prevented us from performing replication analyses for the novel associations, e.g. those for heart failure and venous thromboembolism. The two-sample design also hinders the possibility of conducting more detailed analyses such as subgroup stratification based on lipid levels. This design still allows sensitivity analyses such as MR-Egger and the weighted median estimator which enabled us to investigate and account for the possible presence of genetic pleiotropy, and gene-specific analyses that are less likely to be influenced by pleiotropy as they only include variants from a single gene region where the function is well-known. However, while we have taken steps to mitigate against results being driven by pleiotropy as far as possible, we acknowledge that the Mendelian randomization approach is subject to untestable assumptions, and that these may be violated.

Although the principles of Mendelian randomization seek to emulate a randomized trial, the approach is fundamentally observational in nature. While there has generally been broad concordance between results from Mendelian randomization investigations and findings from randomized trials (as demonstrated by the results shown here for CAD), there are reasons why quantitative results could differ between the approaches.^43^ Genetic variants lead to small but long-term changes in the average levels of risk factors, whereas trials tend to be shorter in duration but assess larger changes in risk factors. Additionally, this investigation was conducted in UK-based middle- to late-aged participants of European ancestries. While this is recommended for Mendelian randomization to ensure that genetic associations are not influenced by population stratification, it means that results may not be generalizable to other ethnicities or nationalities.

In conclusion, multivariable Mendelian randomization analyses accounting for lipid-related genetic pleiotropy support the hypothesis that circulating lipids have causal effects on a wide range of cardiovascular diseases. Interventions that lower LDL-cholesterol are likely to be beneficial in preventing aortic aneurysm and aortic stenosis in addition to several other atheromatous diseases. Triglyceride lowering treatments are likely to be beneficial in preventing coronary artery disease and aortic valve stenosis, but caution is needed due to the possibility of increased risk of venous thromboembolism.

## Supplementary Online Materials

### SUPPLEMENTARY METHODS

#### Study design and data sources

We obtained genetic associations with lipid concentrations (LDL-cholesterol, HDL-cholesterol, and triglycerides) from the Global Lipids Genetics Consortium (GLGC) on up to 188,577 participants of European ancestry.^21^ Genetic associations were estimated with adjustment for age, sex, and genomic principal components within each participating study after inverse rank quantile normalization of lipid concentrations, and then meta-analysed across studies. Inverse rank quantile normalization means that associations are not affected by the skewed distribution of variables, in particular for triglycerides. The triglyceride measurement mostly represents the triglyceride content of triglyceride-rich lipoproteins.

We estimated genetic associations with disease outcomes from UK Biobank, a cohort of 502,682 participants (94% of self-reported European ancestry) recruited between 2006-2010 in 22 assessment centres throughout the UK and followed-up until 31st March 2017 or their date of death (recorded until 14th February 2018).^22^ In addition to standard quality control procedures,^44^ we excluded participants having non-European ancestry (self-report or inferred by genetics), low call rate or excess heterozygosity (>3 standard deviations from the mean). We included only one of each set of related participants (third-degree relatives or closer). The sample for our analyses comprised 367,703 unrelated participants of European ancestry.

We defined 19 outcomes based on electronic health records (ICD-9 or ICD-10 diagnosis and hospital procedure codes) from hospital episode statistics and death certificates, and self-reported information validated by interview at baseline with a trained nurse. With respect to study baseline, we included both prevalent and incident events in our outcome definitions. However, with respect to the genetic variants, all events are incident. For presentation, we divide outcomes into five categories: 1) ischaemic cerebrovascular diseases (ischaemic stroke, transient ischaemic attack, combined ischaemic cerebrovascular event [i.e. ischaemic stroke or transient ischaemic attack]), 2) haemorrhagic stroke (intracerebral haemorrhage, subarachnoid haemorrhage, combined haemorrhagic stroke); 3) aneurysms (any aortic aneurysm, abdominal aortic aneurysm, thoracic aortic aneurysm); 4) thromboembolic diseases (any venous thromboembolism, pulmonary embolism, deep vein thrombosis); and 5) other cardiovascular diseases (hypertension, peripheral vascular disease, aortic valve stenosis, atrial fibrillation, and heart failure). Additionally, we included CAD as a positive control outcome and chronic kidney disease as a negative control outcome as it is thought to be influenced by some cardiovascular risk factors (such as hypertension and diabetes) but not dyslipidaemia.^23^ Precise outcome definitions are given in Supplementary Table S1. To obtain genetic association estimates for each outcome, we conducted logistic regression with adjustment for age at recruitment, sex and 10 genomic principal components using the *snptest* program.^45^

#### Statistical analysis

We carried out analyses based on 184 genetic variants previously demonstrated to be associated with at least one of LDL-cholesterol, HDL-cholesterol or triglycerides at a genome-wide level of significance (*p* < 5 × 10^−8^) in the GLGC. Only one variant per gene region was included in the analysis, except for a small number of regions where a conditionally independent signal was found. We accounted for correlations between variants in the analysis methods, except for the weighted median method, where we further pruned to only include one variant per gene region. These variants explained 13.7% of the variance in HDL-cholesterol, 14.6% in LDL-cholesterol, and 11.7% in triglycerides in GLGC. One variant (rs2954022) was omitted from analyses as it was triallelic in UK Biobank but all other variants were well-imputed (info score >0.9) and were retained.

To obtain the associations of genetically-predicted values of these lipids with each cardiovascular outcome while accounting for measured genetic pleiotropy via other major lipids, we performed multivariable Mendelian randomization analyses.^24^ We implemented this method by weighted regression of the genetic associations with the outcome on the genetic associations with the three predictors, while fixing the intercept to be zero. For sensitivity analysis, we also performed (i) univariable Mendelian randomization using the inverse-variance weighted method for each lipid fraction based on all variants associated with that lipid fraction at a genome-wide level of significance,^25^ (ii) MR-Egger regression to account for unmeasured pleiotropy,^26^ and (iii) weighted median regression to assess robustness to invalid genetic instruments.^27^ To account for between-variant heterogeneity, we used multiplicative random-effects models in all analyses. All Mendelian randomization estimates are expressed per 1 standard deviation increase in the corresponding lipid fraction in GLGC (1 standard deviation was 39.0 mg/dL for LDL-cholesterol, 15.8 mg/dL for HDL-cholesterol, and 90.5 mg/dL for triglycerides).

For cardiovascular diseases for which it is possible that effect of lipid fractions on that outcome may act via other conditions (e.g. via CAD for heart failure), we also performed multivariable Mendelian randomization analyses adjusted for the hypothesised mediating condition. This analysis estimates the direct effects of the lipid fractions on the outcome that are not mediated via CAD. The coefficient for CAD represents the odds ratio for the outcome per unit increase in the beta-coefficient for CAD (i.e. per unit increase in log odds ratio for CAD) Additionally, we performed multivariable Mendelian randomization analyses for these outcomes excluding participants with a CAD diagnosis, to assess whether associations persisted in participants without comorbid CAD. To assess the effect of triglycerides on molecular traits related to thrombosis, we searched publicly-available genetic associations from Phenoscanner (http://www.phenoscanner.medschl.cam.ac.uk/),^28^ a curated database of over 60 billion genetic associations, and performed multivariable Mendelian randomization using these traits as outcomes.

We also performed gene-specific analyses for variants in specific gene regions that can be considered as proxies for existing or proposed lipid-lowering therapies (since the protein products of these genes are the pharmacological targets of these drugs) and that explain enough variance in lipid concentration to result in adequately powered analyses (at least 0.4% for at least one lipid measure). We considered the following gene regions: *HMGCR* (proxy for statin treatment), *PCSK9* (proxy for PCSK9 inhibitors), *LDLR* (proxy for inhibition of the LDL receptor), *APOC3* (proxy for APOC3 inhibitors), and *LPL* (proxy for lipoprotein lipase inhibition). For each gene region, we selected genetic variants that were not strongly correlated (R^2^<0.8) with one another and that were previously associated with LDL-cholesterol in a conditional analysis (Supplementary Table S2). All variants are within 500kb of the gene. We obtained univariable Mendelian randomization estimates using the inverse-variance weighted method while accounting for correlations between variants.^25^ Variants in each gene region explained 0.4% (*HMGCR*), 1.2% (*PCSK9*), 1.0% (*LDLR*), 0.1% (*APOC3*) and less than 0.1% (*LPL*) of the variance in LDL-cholesterol in GLGC. The *APOC3* and *LPL* variants also explained 1.0% and 0.9% of the variance in triglycerides respectively. The power of the gene-specific analyses differs considerably owing to the different proportion of variance explained. Gene-specific estimates are expressed per 1 standard deviation increase in LDL-cholesterol for the *HMGCR, PCSK9*, and *LDLR* regions, and per 1 standard deviation increase in triglycerides for the *APOC3* and *LPL* regions.

We carried out all analyses using R (version 3.4.4) unless otherwise stated. All p-values presented are two-sided.

## SUPPLEMENTARY RESULTS

**Supplementary Table S1:** Summary of disease outcomes considered: sources of information

| OUTCOME | NUMBER OF CASES | ICD-9 DIAGNOSIS | ICD-10 DIAGNOSIS | OPCS PROCEDURE | SELF-REPORT* |
| --- | --- | --- | --- | --- | --- |
| *Coronary Artery Disease* | 29,278 | 410.X, 411.X, 412.X, 414.0, 414.8, 414.9 | I21.X, I22.X, I23.X, I24.X, I25.1, I25.2, I25.5, I25.6, I25.8, I25.9 | K40.X, K41.X, K42.X, K43.X, K44.X, K45.X, K46.X, K49.X, K50.1, K50.2, K50.4, K75.X | Non-cancer illness code (20002), Surgical operation code (20004), Health condition diagnosed by doctor (6150) |
| *Ischaemic Cerebrovascular Disease (all)* | 8084 | 433.X, 434.X, 435.X | G45.X, I63.X |  | Non-cancer illness code (20002) |
| *Ischaemic Stroke* | 4602 | 433.X, 434.X | I63.X |  | Non-cancer illness code (20002) |
| *Transient Ischaemic Attack* | 3962 | 435.X | G45.X |  | Non-cancer illness code (20002) |
| *Haemorrhagic Stroke (all)* | 1981 | 430.X, 431.X | I60.X, I61.X |  | Non-cancer illness code (20002) |
| *Intracerebral Haemorrhage* | 1064 | 431.X | I61.X |  | Non-cancer illness code (20002) |
| *Subarachnoid Haemorrhage* | 1084 | 430.X | I60.X |  | Non-cancer illness code (20002) |
| *Aortic Aneurysm (all)* | 1849 | 441.X | I71.X | L19.4, L19.5 | Non-cancer illness code (20002), Surgical operation code (20004) |
| *Abdominal Aortic Aneurysm* | 1094 | 441.3, 441.4 | I71.3, I71.4 | L19.4, L19.5 | Non-cancer illness code (20002) |
| *Thoracic Aortic Aneurysm* | 347 | 441.1, 441.2 | I71.1, I71.2 |  | Non-cancer illness code (20002) |
| *Venous Thromboembolism (all)* | 14,097 | 415.1, 451.1, 452.X, 453.0, 453.4, 453.9, | I26.X, I80.1, I80.2, I81.X, I82.0 | L90.2 | Non-cancer illness code (20002), Health condition diagnosed by doctor (6152) |
| *Deep Vein Thrombosis* | 9454 | 451.1 | I80.2 | L90.2 | Non-cancer illness code (20002), Health condition diagnosed by doctor (6152) |
| *Pulmonary Embolism* | 6148 | 415.1 | I26.X |  | Non-cancer illness code (20002), Health condition diagnosed by doctor (6152) |
| *Hypertension* | 125,846 | 401.X | I10 |  | Non-cancer illness code (20002), Health condition diagnosed by doctor (6150), Medication for health condition (6177) |
| *Peripheral Vascular Disease* | 3415 | 443.8, 443.9 | I73.8, I73.9 |  | Non-cancer illness code (20002) |
| *Aortic Valve Stenosis* | 2244 |  | I35.0, I35.2 |  | Non-cancer illness code (20002) |
| *Atrial Fibrillation* | 16,945 | 427.3 | I48 |  | Non-cancer illness code (20002) |
| *Heart Failure* | 6712 | 402.01, 402.11, 402.91, 404.01, 404.11, 404.91, 404.03, 404.13, 404.93, 428.X | I11.0, I13.0, I13.2, I50.X |  | Non-cancer illness code (20002) |
| *Chronic Kidney Disease* | 6321 | 585.X | N18.X |  | Non-cancer illness code (20002) |

Note that .X means that all sub-codes are matched. Abbreviations: ICD = International Classification of Disease; OPCS = Office of Population Censuses and Surveys Classification of Surgical Operations and Procedures; TIA = transient ischaemic attack
\* Health condition diagnosed by doctor (6150/6152) and Medication for health condition (6177) were self-reported from touchscreen; Non-cancer illness code (20002) and Surgical operation code (20004) were self-reported from interview with trained nurse.

**Supplementary Table S2:** Genetic variants included in gene-specific analyses for each region.

| GENE<br>REGION | SNP | POS (HG19) | EFFECT<br>ALLELE | OTHER<br>ALLELE | EFFECT<br>ALLELE<br>FREQUENCY | BETA | SE | P-VALUE |
| --- | --- | --- | --- | --- | --- | --- | --- | --- |
| *HMGCR* | rs2006760 | chr5:74562029 | G | C | 0.229 | 0.053 | 0.008 | $1.7 \times 10^{-13}$ |
| *HMGCR* | rs2303152 | chr5: 74641707 | G | A | 0.890 | -0.042 | 0.006 | $1.1 \times 10^{-9}$ |
| *HMGCR* | rs17238484 | chr5:74648496 | G | T | 0.763 | -0.063 | 0.006 | $1.4 \times 10^{-21}$ |
| *HMGCR* | rs12916 | chr5:74656539 | C | T | 0.412 | 0.073 | 0.004 | $7.8 \times 10^{-78}$ |
| *HMGCR* | rs10066707 | chr5: 74560579 | A | G | 0.396 | 0.050 | 0.005 | $3.0 \times 10^{-19}$ |
| *HMGCR* | rs5909 | chr5: 74656175 | A | G | 0.100 | 0.062 | 0.009 | $5.0 \times 10^{-13}$ |
| *PCSK9* | rs2479394 | chr1:55486064 | G | A | 0.281 | 0.039 | 0.004 | $1.6 \times 10^{-19}$ |
| *PCSK9* | rs11206510 | chr1:55496039 | C | T | 0.172 | -0.083 | 0.005 | $2.4 \times 10^{-53}$ |
| *PCSK9* | rs2149041 | chr1:55502137 | C | G | 0.823 | -0.064 | 0.005 | $1.4 \times 10^{-35}$ |
| *PCSK9* | rs10888897 | chr1:55513061 | C | T | 0.680 | -0.064 | 0.004 | $2.5 \times 10^{-50}$ |
| *PCSK9* | rs7552841 | chr1:55518752 | C | T | 0.596 | 0.051 | 0.004 | $8.4 \times 10^{-31}$ |
| *PCSK9* | rs562556 | chr1:55524237 | A | G | 0.631 | -0.037 | 0.004 | $5.4 \times 10^{-15}$ |
| *LDLR* | rs6511720 | chr19:11202306 | G | T | 0.902 | 0.221 | 0.006 | $3.9 \times 10^{-262}$ |
| *LDLR* | rs688 | chr19:11227602 | C | T | 0.553 | -0.054 | 0.004 | $1.0 \times 10^{-43}$ |
| *APOC3* | rs10790162 | chr11:116639104 | A | G | 0.091 | 0.231 | 0.006 | $1.1 \times 10^{-249}$ |
| *APOC3* | rs603446 | chr11:116654435 | C | T | 0.553 | 0.050 | 0.003 | $3.9 \times 10^{-43}$ |
| *LPL* | rs1801177 | chr8:19805708 | A | G | 0.011 | 0.164 | 0.023 | $1.1 \times 10^{-9}$ |
| *LPL* | rs268 | chr8:19813529 | A | G | 0.986 | -0.280 | 0.035 | $2.2 \times 10^{-16}$ |
| *LPL* | rs301 | chr8:19816934 | C | T | 0.459 | -0.109 | 0.004 | $1.9 \times 10^{-167}$ |
| *LPL* | rs326 | chr8:19819439 | A | G | 0.673 | 0.087 | 0.005 | $1.0 \times 10^{-63}$ |
| *LPL* | rs328 | chr8:19819724 | C | G | 0.870 | 0.167 | 0.006 | $2.0 \times 10^{-179}$ |

Genetic associations (beta and standard error [SE]) are with LDL-cholesterol for the *HMGCR*, *PCSK9*, and *LDLR* gene regions, and with triglycerides for the *APOC3* and *LPL* gene regions. They are given in standard deviation units and are taken from the Global Lipids Genetics Consortium (see http://csg.sph.umich.edu/willer/public/lipids2013/).

**Supplementary Table S3:** Estimates (odds ratio per 1 standard deviation increase in lipid fraction and 95% confidence interval) from polygenic multivariable Mendelian randomization analyses for all lipid-related variants. Estimates with *p* < 0.05 are reported in **bold**.

| DISEASE | HDL-CHOLESTEROL | LDL-CHOLESTEROL | TRIGLYCERIDES |
| --- | --- | --- | --- |
| *Coronary Artery Disease* | **0.91 (0.83-1.00)** | **1.45 (1.35-1.57)** | **1.25 (1.12-1.40)** |
| *Ischaemic Cerebrovascular Disease (all)* | 0.95 (0.86-1.05) | **1.14 (1.04-1.24)** | 0.98 (0.87-1.10) |
| *Ischaemic Stroke* | 0.98 (0.86-1.12) | 1.10 (0.98-1.23) | 1.03 (0.87-1.20) |
| *Transient Ischaemic Attack* | 0.91 (0.80-1.03) | **1.16 (1.04-1.29)** | 0.94 (0.81-1.10) |
| *Haemorrhagic Stroke (all)* | **0.76 (0.63-0.92)** | 0.90 (0.76-1.05) | **0.78 (0.62-0.98)** |
| *Intracerebral Haemorrhage* | **0.65 (0.51-0.82)** | 0.86 (0.70-1.05) | **0.65 (0.49-0.86)** |
| *Subarachnoid Haemorrhage* | 0.82 (0.63-1.06) | 0.90 (0.72-1.12) | 0.85 (0.62-1.17) |
| *Aortic Aneurysm (all)* | **0.79 (0.65-0.96)** | **1.43 (1.21-1.68)** | 1.06 (0.84-1.33) |
| *Abdominal Aortic Aneurysm* | **0.65 (0.51-0.85)** | **1.75 (1.40-2.17)** | 0.94 (0.68-1.28) |
| *Thoracic Aortic Aneurysm* | 0.81 (0.53-1.22) | 0.98 (0.69-1.38) | 0.93 (0.57-1.54) |
| *Venous Thromboembolism (all)* | 0.94 (0.82-1.08) | 1.03 (0.92-1.16) | **0.79 (0.67-0.93)** |
| *Deep Vein Thrombosis* | 0.94 (0.81-1.09) | 1.00 (0.88-1.13) | **0.78 (0.65-0.93)** |
| *Pulmonary Embolism* | 0.98 (0.83-1.17) | 1.12 (0.97-1.29) | **0.78 (0.64-0.96)** |
| *Hypertension* | **0.90 (0.83-0.97)** | 1.05 (0.98-1.12) | **1.17 (1.07-1.27)** |
| *Peripheral Vascular Disease* | 0.88 (0.75-1.03) | **1.15 (1.01-1.31)** | 1.01 (0.84-1.22) |
| *Aortic Valve Stenosis* | 0.98 (0.82-1.18) | **1.46 (1.25-1.70)** | **1.29 (1.04-1.61)** |
| *Atrial Fibrillation* | 0.94 (0.87-1.03) | 1.07 (1.00-1.15) | 0.93 (0.84-1.02) |
| *Heart Failure* | 0.97 (0.87-1.08) | **1.17 (1.06-1.28)** | **1.19 (1.04-1.37)** |
| *Chronic Kidney Disease* | 0.90 (0.81-1.00) | 0.93 (0.85-1.02) | 0.98 (0.87-1.12) |

**Supplementary Table S4:** Associations of genetically-predicted triglyceride levels with traits relating to thrombosis and coagulation.

| TRAIT | STUDY | PARTICIPANTS | VARIANTS | BETA | P-VALUE | DESCRIPTION | LIKELY ROLE |
| --- | --- | --- | --- | --- | --- | --- | --- |
| *Metalloproteinase Inhibitor 4* | Sun 2017<sup>46</sup> | 3,301 | 181 | -0.378 | <0.001 | Metalloproteinase inhibition reduces thrombolytic (tissue plasminogen activator)-induced cerebral haemorrhage<sup>47</sup> | Pro-thrombotic |
| *Red Cell Distribution Width* | Astle 2016<sup>48</sup> | 173,480 | 182 | -0.169 | <0.001 | Positively associated with risk of venous thromboembolism<sup>49, 50</sup> | Pro-thrombotic |
| *Matrix Metalloproteinase-9* | Sun 2017<sup>46</sup> | 3,301 | 181 | 0.256 | 0.002 | Found in the vein wall during thrombus resolution<sup>51</sup> | Anti-thrombotic |
| *Plasminogen Activator Inhibitor 1* | Sun 2017<sup>46</sup> | 3,301 | 181 | 0.244 | 0.007 | Plasminogen activator inhibitor 1 can suppress the generation of active plasmin which eliminates fibrin<sup>52</sup> and has been positively associated with venous thromboembolism<sup>53</sup> | Pro-thrombotic |
| *Kallikrein-4* | Sun 2017<sup>46</sup> | 3,301 | 181 | 0.212 | 0.011 | Kallikrein-deficient mice leads to suppression of vessel wall tissue factor, which leads to reduced thrombosis<sup>54</sup> | Anti-thrombotic |
| *Tissue-type Plasminogen Activator* | Folkersen 2017<sup>55</sup> | 3,394 | 179 | 0.185 | 0.031 | Catalyses conversion of plasminogen to plasmin, thus promoting fibrinolysis<sup>56</sup> | Anti-thrombotic |
| *Coagulation Factor Xa* | Sun 2017<sup>46</sup> | 3,301 | 181 | 0.186 | 0.034 | Protease involved in the common coagulation pathway | Pro-thrombotic |

Beta coefficients are scaled to a standard deviation (SD) change in the trait per 1 SD increase in genetically-predicted triglyceride levels. Estimates were obtained from multivariable Mendelian randomization accounting for genetic associations with LDL-cholesterol and HDL-cholesterol.

**Supplementary Figure S1:**
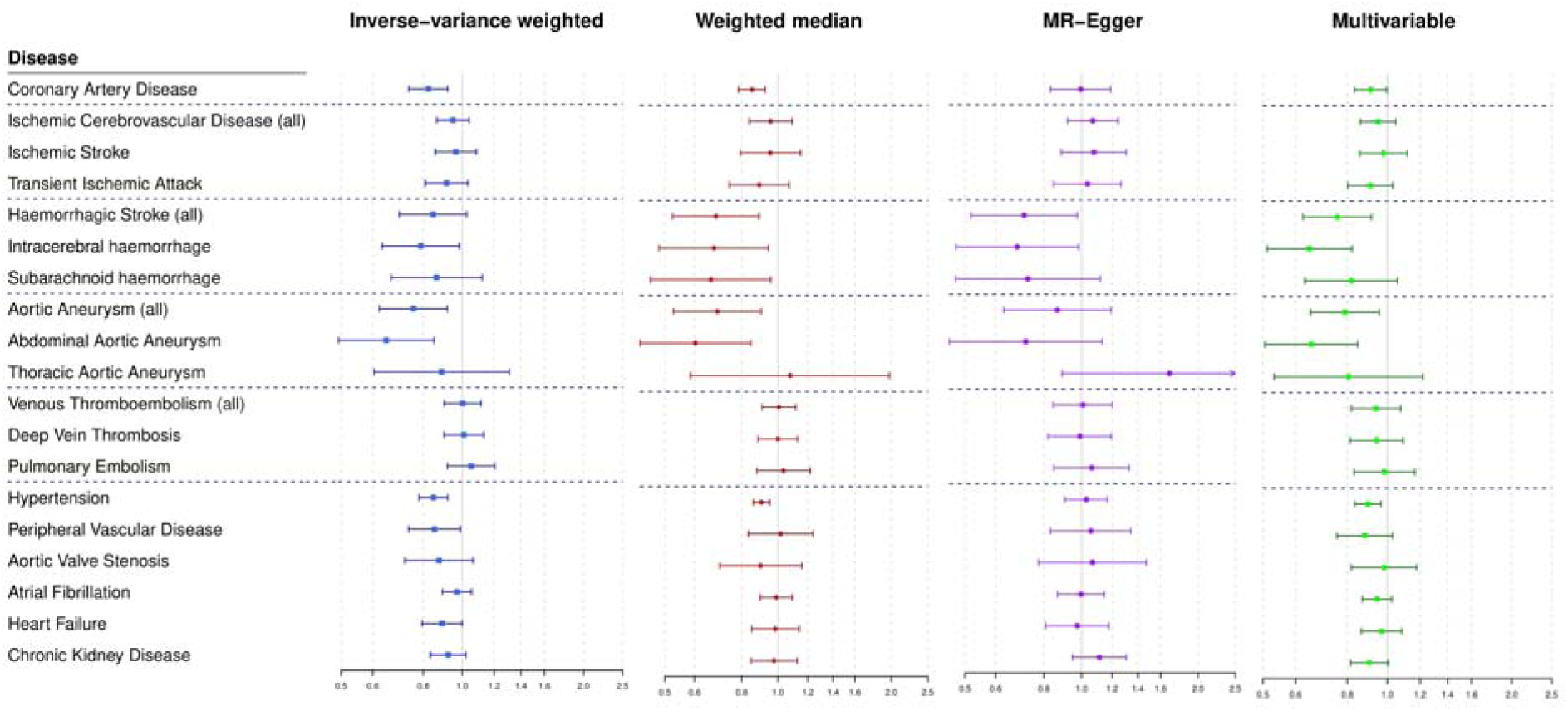
Estimates (odds ratio per 1 standard deviation increase in lipid fraction and 95% confidence interval) for HDL-cholesterol from genome-wide Mendelian randomization analyses for all lipid-related variants associated with the target lipid fraction.

**Supplementary Table S5:** Estimates (odds ratio per 1 standard deviation increase in lipid fraction and 95% confidence interval) for HDL-cholesterol from polygenic univariable Mendelian randomization analyses for all lipid-related variants associated with the target lipid fraction. Estimates with *p* < 0.05 are reported in **bold**.

| DISEASE | INVERSE-VARIANCE WEIGHTED | WEIGHTED MEDIAN | MR-EGGER REGRESSION |
| --- | --- | --- | --- |
| *Coronary Artery Disease* | **0.82 (0.74-0.92)** | **0.85 (0.78-0.92)** | 0.99 (0.83-1.19) |
| *Ischaemic Cerebrovascular Disease (all)* | 0.95 (0.86-1.04) | 0.96 (0.84-1.09) | 1.07 (0.92-1.24) |
| *Ischaemic Stroke* | 0.96 (0.86-1.08) | 0.95 (0.79-1.15) | 1.08 (0.89-1.31) |
| *Transient Ischaemic Attack* | 0.91 (0.81-1.03) | 0.89 (0.74-1.07) | 1.04 (0.85-1.27) |
| *Haemorrhagic Stroke (all)* | 0.85 (0.70-1.03) | **0.68 (0.52-0.89)** | **0.71 (0.52-0.97)** |
| *Intracerebral Haemorrhage* | **0.79 (0.63-0.98)** | **0.68 (0.48-0.94)** | **0.68 (0.47-0.98)** |
| *Subarachnoid Haemorrhage* | 0.86 (0.67-1.12) | **0.66 (0.46-0.96)** | 0.73 (0.47-1.12) |
| *Aortic Aneurysm (all)* | **0.76 (0.62-0.92)** | **0.69 (0.53-0.90)** | 0.87 (0.63-1.19) |
| *Abdominal Aortic Aneurysm* | **0.65 (0.49-0.85)** | **0.60 (0.43-0.85)** | 0.72 (0.46-1.13) |
| *Thoracic Aortic Aneurysm* | 0.89 (0.60-1.31) | **1.08 (0.59-1.98)** | 1.69 (0.89-3.19) |
| *Venous Thromboembolism (all)* | 1.00 (0.90-1.11) | 1.01 (0.91-1.11) | 1.01 (0.85-1.20) |
| *Deep Vein Thrombosis* | 1.01 (0.90-1.13) | 1.00 (0.88-1.13) | 0.99 (0.82-1.20) |
| *Pulmonary Embolism* | 1.05 (0.92-1.20) | 1.03 (0.88-1.22) | 1.06 (0.85-1.33) |
| *Hypertension* | **0.85 (0.78-0.92)** | **0.90 (0.86-0.95)** | 1.03 (0.91-1.17) |
| *Peripheral Vascular Disease* | **0.85 (0.74-0.99)** | 1.02 (0.83-1.24) | 1.06 (0.83-1.34) |
| *Aortic Valve Stenosis* | 0.88 (0.72-1.06) | 0.90 (0.70-1.16) | 1.07 (0.77-1.47) |
| *Atrial Fibrillation* | 0.97 (0.89-1.05) | 0.99 (0.90-1.09) | 1.00 (0.87-1.15) |
| *Heart Failure* | **0.89 (0.79-1.00)** | 0.98 (0.85-1.14) | 0.97 (0.81-1.18) |
| *Chronic Kidney Disease* | 0.92 (0.83-1.02) | 0.98 (0.85-1.13) | 1.11 (0.95-1.31) |

**Supplementary Figure S2:**
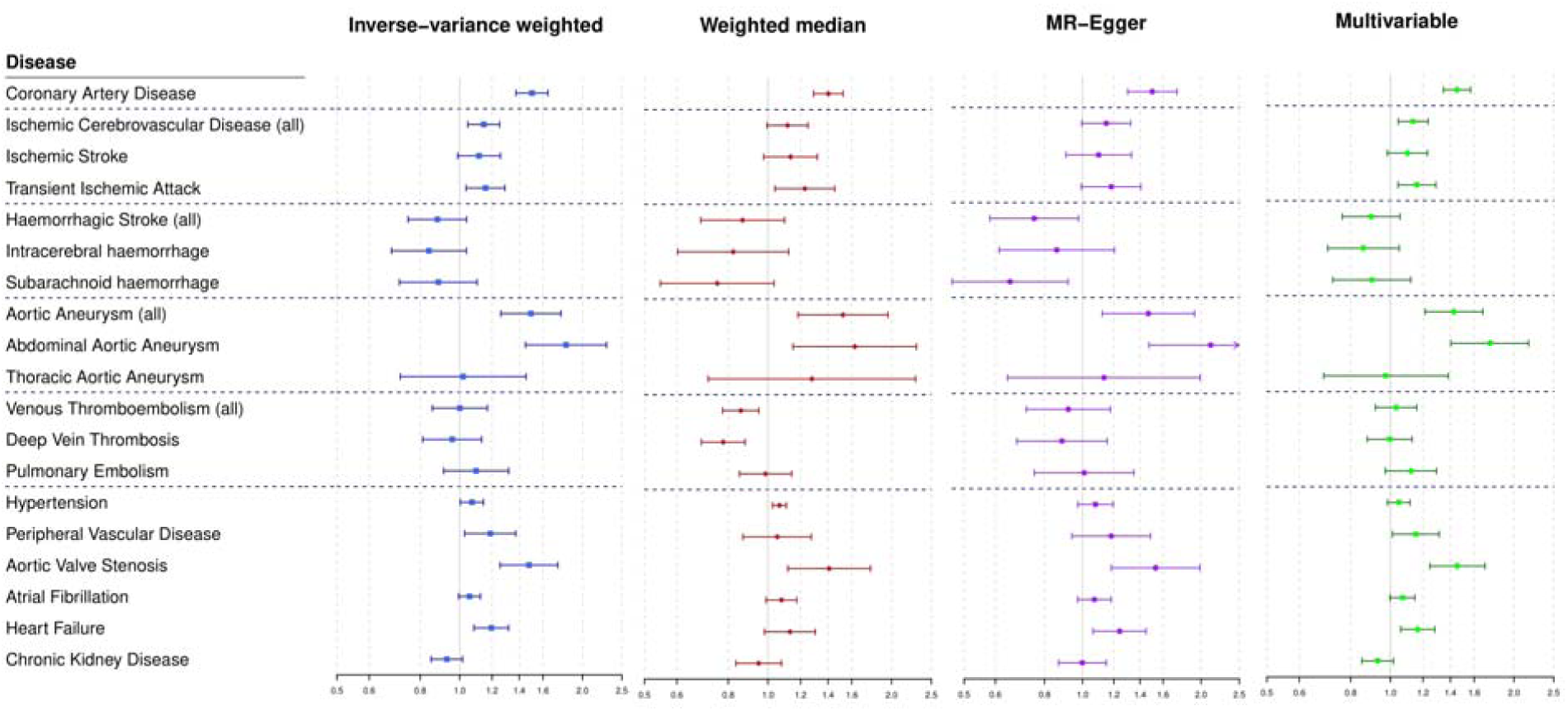
Estimates (odds ratio per 1 standard deviation increase in lipid fraction and 95% confidence interval) for LDL-cholesterol from genome-wide Mendelian randomization analyses for all lipid-related variants associated with the target lipid fraction.

**Supplementary Table S6:** Estimates (odds ratio per 1 standard deviation increase in lipid fraction and 95% confidence interval) for LDL-cholesterol from polygenic univariable Mendelian randomization analyses for all lipid-related variants associated with the target lipid fraction. Estimates with *p* < 0.05 are reported in **bold**.

| DISEASE | INVERSE-<br>VARIANCE<br>WEIGHTED | WEIGHTED<br>MEDIAN | MR-EGGER<br>REGRESSION |
| --- | --- | --- | --- |
| *Coronary Artery Disease* | **1.50 (1.37-1.65)** | **1.40 (1.29-1.52)** | **1.50 (1.30-1.74)** |
| *Ischaemic Cerebrovascular Disease (all)* | **1.15 (1.05-1.25)** | 1.12 (0.99-1.25) | 1.15 (1.00-1.32) |
| *Ischaemic Stroke* | 1.12 (0.99-1.26) | 1.13 (0.98-1.32) | 1.10 (0.91-1.33) |
| *Transient Ischaemic Attack* | **1.16 (1.04-1.29)** | **1.23 (1.04-1.45)** | 1.18 (0.99-1.41) |
| *Haemorrhagic Stroke (all)* | 0.88 (0.75-1.04) | 0.87 (0.69-1.10) | **0.75 (0.58-0.98)** |
| *Intracerebral Haemorrhage* | 0.84 (0.68-1.04) | 0.82 (0.60-1.12) | 0.86 (0.61-1.20) |
| *Subarachnoid Haemorrhage* | 0.89 (0.71-1.10) | 0.75 (0.55-1.03) | **0.65 (0.47-0.92)** |
| *Aortic Aneurysm (all)* | **1.50 (1.26-1.77)** | **1.52 (1.18-1.96)** | **1.47 (1.12-1.92)** |
| *Abdominal Aortic Aneurysm* | **1.82 (1.45-2.29)** | **1.63 (1.15-2.30)** | **2.12 (1.48-3.04)** |
| *Thoracic Aortic Aneurysm* | 1.02 (0.71-1.46) | 1.28 (0.71-2.29) | 1.13 (0.64-1.99) |
| *Venous Thromboembolism (all)* | 1.00 (0.86-1.17) | **0.86 (0.78-0.95)** | 0.92 (0.72-1.18) |
| *Deep Vein Thrombosis* | 0.96 (0.81-1.13) | **0.78 (0.69-0.88)** | 0.89 (0.68-1.15) |
| *Pulmonary Embolism* | 1.10 (0.91-1.32) | 0.99 (0.85-1.14) | 1.01 (0.75-1.35) |
| *Hypertension* | **1.07 (1.00-1.14)** | **1.07 (1.03-1.11)** | 1.08 (0.97-1.20) |
| *Peripheral Vascular Disease* | **1.19 (1.03-1.37)** | 1.05 (0.87-1.27) | 1.18 (0.94-1.49) |
| *Aortic Valve Stenosis* | **1.48 (1.26-1.74)** | **1.41 (1.12-1.78)** | **1.53 (1.18-1.99)** |
| *Atrial Fibrillation* | 1.06 (0.99-1.12) | 1.08 (0.99-1.17) | 1.07 (0.97-1.18) |
| *Heart Failure* | **1.19 (1.08-1.32)** | 1.13 (0.98-1.30) | **1.24 (1.06-1.45)** |
| *Chronic Kidney Disease* | 0.93 (0.85-1.02) | 0.95 (0.84-1.08) | 1.00 (0.87-1.15) |

**Supplementary Figure S3:**
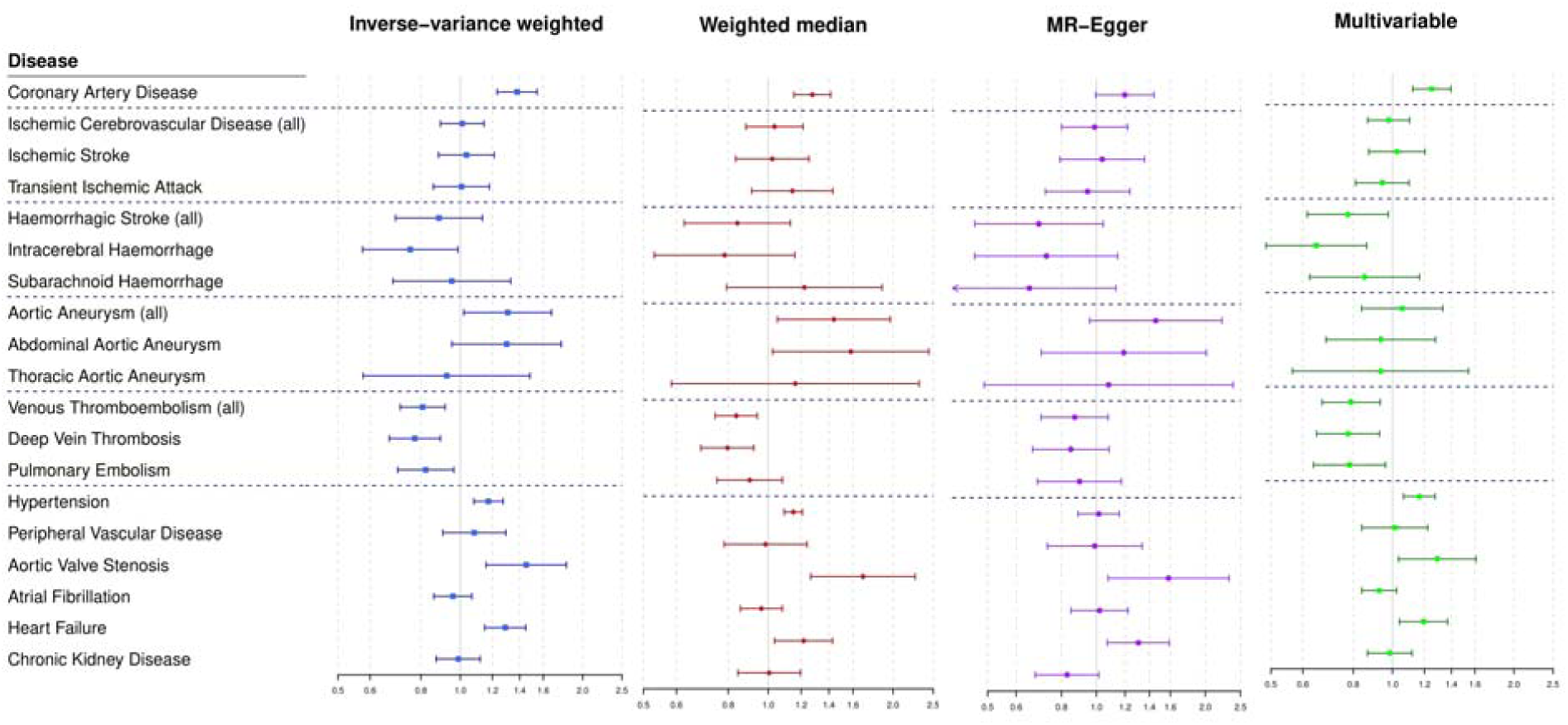
Estimates (odds ratio per 1 standard deviation increase in lipid fraction and 95% confidence interval) for triglycerides from genome-wide Mendelian randomization analyses for all lipid-related variants associated with the target lipid fraction.

**Supplementary Table S7:** Estimates (odds ratio per 1 standard deviation increase in lipid fraction and 95% confidence interval) for triglycerides from polygenic univariable Mendelian randomization analyses for all lipid-related variants associated with the target lipid fraction. Estimates with *p* < 0.05 are reported in **bold**.

| DISEASE | INVERSE-VARIANCE WEIGHTED | WEIGHTED MEDIAN | MR-EGGER REGRESSION |
| --- | --- | --- | --- |
| *Coronary Artery Disease* | **1.38 (1.23-1.54)** | **1.28 (1.15-1.41)** | 1.20 (0.99-1.44) |
| *Ischaemic Cerebrovascular Disease (all)* | 1.01 (0.89-1.14) | 1.04 (0.88-1.21) | 0.99 (0.80-1.22) |
| *Ischaemic Stroke* | 1.03 (0.88-1.21) | 1.02 (0.83-1.25) | 1.04 (0.79-1.36) |
| *Transient Ischaemic Attack* | 1.01 (0.86-1.18) | 1.14 (0.91-1.43) | 0.94 (0.72-1.24) |
| *Haemorrhagic Stroke (all)* | 0.89 (0.69-1.13) | 0.84 (0.63-1.13) | 0.69 (0.46-1.04) |
| *Intracerebral Haemorrhage* | **0.75 (0.57-0.99)** | 0.78 (0.53-1.16) | 0.73 (0.46-1.15) |
| *Subarachnoid Haemorrhage* | 0.95 (0.68-1.33) | 1.22 (0.79-1.88) | 0.65 (0.38-1.13) |
| *Aortic Aneurysm (all)* | **1.31 (1.02-1.68)** | **1.44 (1.05-1.97)** | 1.46 (0.96-2.22) |
| *Abdominal Aortic Aneurysm* | 1.30 (0.95-1.77) | **1.58 (1.02-2.44)** | 1.19 (0.70-2.01) |
| *Thoracic Aortic Aneurysm* | 0.93 (0.58-1.48) | 1.16 (0.58-2.31) | 1.08 (0.49-2.39) |
| *Venous Thromboembolism (all)* | **0.81 (0.71-0.92)** | **0.84 (0.74-0.94)** | 0.87 (0.70-1.08) |
| *Deep Vein Thrombosis* | **0.77 (0.67-0.89)** | **0.80 (0.69-0.92)** | 0.85 (0.67-1.08) |
| *Pulmonary Embolism* | **0.82 (0.70-0.96)** | 0.90 (0.75-1.08) | 0.90 (0.69-1.17) |
| *Hypertension* | **1.17 (1.08-1.27)** | **1.15 (1.09-1.21)** | 1.01 (0.89-1.16) |
| *Peripheral Vascular Disease* | 1.08 (0.91-1.29) | 0.98 (0.78-1.24) | 0.99 (0.73-1.34) |
| *Aortic Valve Stenosis* | **1.45 (1.16-1.82)** | **1.69 (1.27-2.26)** | **1.58 (1.08-2.32)** |
| *Atrial Fibrillation* | 0.96 (0.86-1.07) | 0.96 (0.86-1.08) | 1.02 (0.85-1.22) |
| *Heart Failure* | **1.29 (1.15-1.45)** | **1.22 (1.04-1.43)** | **1.30 (1.07-1.59)** |
| *Chronic Kidney Disease* | 0.99 (0.87-1.12) | 1.00 (0.84-1.20) | 0.83 (0.68-1.01) |

**Supplementary Table S8:** Estimates (odds ratio per 1 standard deviation increase in lipid fraction and 95% confidence interval) from polygenic multivariable Mendelian randomization analyses for all lipid-related variants for heart failure in participants without a CAD diagnosis.

|  | **HDL-cholesterol** | **LDL-cholesterol** | **Triglycerides** |
| --- | --- | --- | --- |
| ***Heart Failure*** | 1.04 (0.88-1.23) | 0.87 (0.75-1.00) | 1.12 (0.91-1.37) |

**Supplementary Table S9:** Multivariable Mendelian randomization estimates (odds ratio with 95% confidence interval) for hypertension, peripheral vascular disease and abdominal aortic aneurysm without and with adjustment for body mass index (BMI) and type 2 diabetes (T2D).

| **Hypertension** | **HDL-cholesterol** | **LDL-cholesterol** | **Triglycerides** | **BMI/T2D** |
| --- | --- | --- | --- | --- |
| *Without adjustment* | 0.90<br>(0.83-0.97)* | 1.05<br>(0.98-1.12) | 1.17<br>(1.07-1.27)* | - |
| *Adjustment for BMI* | 0.91<br>(0.84-0.98)* | 1.06<br>(0.99-1.13) | 1.17<br>(1.07-1.28)* | 1.334<br>(1.08, 1.64)* |
| *Adjustment for T2D* | 0.95<br>(0.89-1.02) | 1.07<br>(1.01-1.13)* | 1.15<br>(1.06-1.25)* | 1.30<br>(1.19-1.40)** |
| **Abdominal aortic aneurysm** | **HDL-cholesterol** | **LDL-cholesterol** | **Triglycerides** | **T2D/BMI** |
| *Without adjustment* | 0.65<br>(0.51-0.85)** | 1.75<br>(1.40-2.17)** | 0.94<br>(0.68-1.28) | - |
| *Adjustment for BMI* | 0.64<br>(0.50-0.83)** | 1.71<br>(1.38-2.13)** | 0.93<br>(0.68-1.27) | 0.57<br>(0.27, 1.21) |
| *Adjustment for T2D* | 0.68<br>(0.52-0.89)** | 1.77<br>(1.42-2.20)** | 0.93<br>(0.68-1.27) | 1.18<br>(0.86-1.61) |
| **Haemorrhagic stroke** | **HDL-cholesterol** | **LDL-cholesterol** | **Triglycerides** | **T2D/BMI** |
| *Without adjustment* | 0.76<br>(0.63-0.92)** | 0.90<br>(0.76-1.05) | 0.78<br>(0.62-0.98)* |  |
| *Adjustment for BMI* | 0.76<br>(0.63-0.92)** | 0.90<br>(0.76-1.06) | 0.77<br>(0.61-0.98)* | 0.98<br>(0.56, 1.70) |
| *Adjustment for T2D* | 0.74<br>(0.61-0.90)** | 0.89<br>(0.76-1.05) | 0.78<br>(0.62-0.98)* | 0.91<br>(0.73-1.15) |

\*: $p < 0.05$ \*\*: $p < 0.005$ \*\*\*: $p < 10^{-8}$
The BMI/T2D column indicates the odds ratio representing the association between genetically-predicted body mass index (per SD increase) or type 2 diabetes (per unit increase in log odds ratio) and the outcomes of interest (i.e. hypertension, abdominal aortic aneurysm, or haemorrhagic stroke) while accounting for the genetic associations of the lipid fractions in a multivariable analysis. If the regression coefficients for the lipid measurements remain unchanged on adjustment for BMI/T2D, then the effects of lipids on the outcome do not operate via BMI/T2D. If the regression coefficients attenuate to the null, then the effects of lipids on the outcome are entirely mediated via BMI/T2D.

**Supplementary Table S10:**
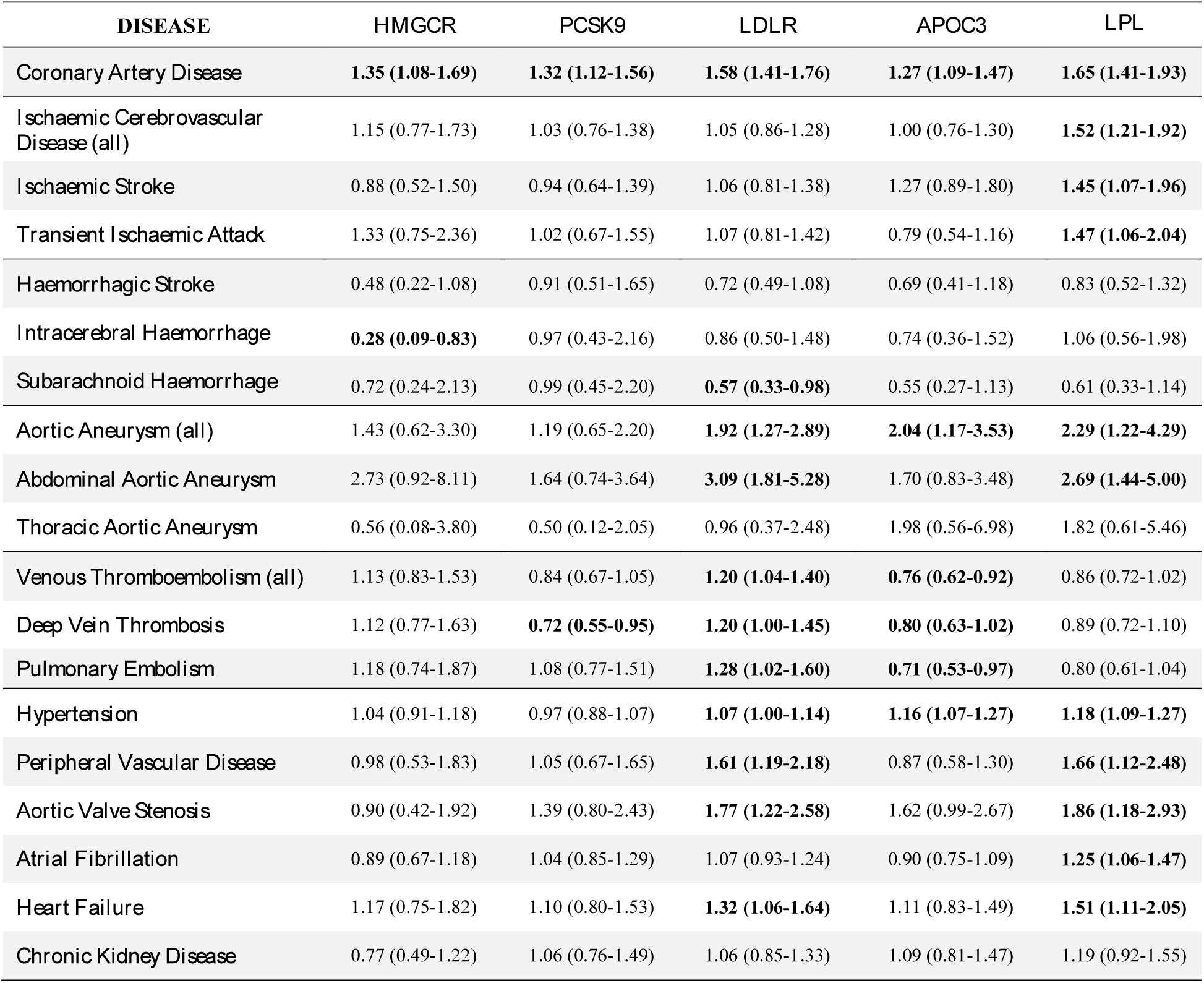
Estimates (odds ratio per 1 standard deviation increase in lipid fraction and 95% confidence interval) from gene-specific Mendelian randomization analyses for four specified gene regions. Estimates with *p* < 0.05 are reported in **bold**. Estimates are calibrated per standard deviation increase in LDL-cholesterol for *HMGCR, PCSK9*, and *LDLR* regions, and per standard deviation increase in triglycerides for *APOC3* and *LPL* regions.

**Supplementary Table S11:** P-values from gene-specific Mendelian randomization analyses for five specified gene regions.

| DISEASE | *HMGCR* | *PCSK9* | *LDLR* | *APOC3* | *LPL* |
| --- | --- | --- | --- | --- | --- |
| *Coronary Artery Disease* | 0.010 | <0.001 | <0.001 | 0.002 | <0.001 |
| *Ischaemic Cerebrovascular Disease (all)* | 0.494 | 0.867 | 0.646 | 0.976 | <0.001 |
| *Ischaemic Stroke* | 0.638 | 0.767 | 0.672 | 0.187 | 0.018 |
| *Transient Ischaemic Attack* | 0.331 | 0.935 | 0.620 | 0.231 | 0.021 |
| *Haemorrhagic Stroke* | 0.076 | 0.762 | 0.109 | 0.175 | 0.427 |
| *Intracerebral Haemorrhage* | 0.022 | 0.934 | 0.596 | 0.415 | 0.864 |
| *Subarachnoid Haemorrhage* | 0.547 | 0.983 | 0.04 | 0.104 | 0.120 |
| *Aortic Aneurysm (all)* | 0.403 | 0.574 | 0.002 | 0.011 | 0.010 |
| *Abdominal Aortic Aneurysm* | 0.070 | 0.223 | <0.001 | 0.144 | 0.002 |
| *Thoracic Aortic Aneurysm* | 0.550 | 0.338 | 0.938 | 0.290 | 0.284 |
| *Venous Thromboembolism (all)* | 0.447 | 0.121 | 0.016 | 0.007 | 0.089 |
| *Deep Vein Thrombosis* | 0.539 | 0.019 | 0.046 | 0.078 | 0.278 |
| *Pulmonary Embolism* | 0.490 | 0.669 | 0.034 | 0.029 | 0.092 |
| *Hypertension* | 0.569 | 0.507 | 0.044 | <0.001 | <0.001 |
| *Peripheral Vascular Disease* | 0.961 | 0.834 | 0.002 | 0.490 | 0.012 |
| *Aortic Valve Stenosis* | 0.782 | 0.244 | 0.003 | 0.057 | 0.008 |
| *Atrial Fibrillation* | 0.409 | 0.690 | 0.324 | 0.295 | 0.008 |
| *Heart Failure* | 0.495 | 0.553 | 0.013 | 0.479 | 0.009 |
| *Chronic Kidney Disease* | 0.271 | 0.719 | 0.597 | 0.586 | 0.189 |

